# Automatic Identification of SARS Coronavirus using Compression-Complexity Measures

**DOI:** 10.1101/2020.03.24.006007

**Authors:** Karthi Balasubramanian, Nithin Nagaraj

## Abstract

Finding vaccine or specific antiviral treatment for global pandemic of virus diseases (such as the ongoing COVID-19) requires rapid analysis, annotation and evaluation of metagenomic libraries to enable a quick and efficient screening of nucleotide sequences. Traditional sequence alignment methods are not suitable and there is a need for fast alignment-free techniques for sequence analysis. Information theory and data compression algorithms provide a rich set of mathematical and computational tools to capture essential patterns in biological sequences. In 2013, our research group (Nagaraj et al., Eur. Phys. J. Special Topics 222(3-4), 2013) has proposed a novel measure known as Effort-To-Compress (ETC) based on the notion of compression-complexity to capture the information content of sequences. In this study, we propose a compression-complexity based distance measure for automatic identification of SARS coronavirus strains from a set of viruses using only short fragments of nucleotide sequences. We also demonstrate that our proposed method can correctly distinguish SARS-CoV-2 from SARS-CoV-1 viruses by analyzing very short segments of nucleotide sequences. This work could be extended further to enable medical practitioners in automatically identifying and characterizing SARS coronavirus strain in a fast and efficient fashion using short and/or incomplete segments of nucleotide sequences. Potentially, the need for sequence assembly can be circumvented.

**Note:** The main ideas and results of this research were first presented at the *International Conference on Nonlinear Systems and Dynamics* (**CNSD-2013**) held at Indian Institute of Technology, Indore, December 12, 2013. In this manuscript, we have extended our preliminary analysis to include SARS-CoV-2 virus as well.

## 1 Introduction

SARS (Severe Acute Respiratory Syndrome) is a viral respiratory disease caused by the SARS coronavirus (SARS-CoV^1^) and having flu-like symptoms. It was first identified in Guandong province, China, in 2002 and spread rapidly to different parts of the world in a span of just a few months [1]. The primary route of transmission of the SARS coronavirus is through mucosal contact with respiratory droplets or fomites of infected persons. Marra *et al.* [1] and Rota *et al.* [2] have done extensive studies to show that SARS-CoVs forms a separate group of coronaviruses and are not closely related to other previously sequenced coronaviruses (mammalian and avian viruses).

SARS-CoV-2 is the latest strain of the coronavirus, first discovered in late 2019 in Wuhan, China, that is responsible for the ongoing pandemic of coronavirus disease 2019 (COVID-19). Apart from SARS-CoV-1 and SARS-CoV-2, there are hundreds of other strains of SARSr-CoV (Severe Acute Respiratory Syndrome-related coronavirus) that are known to infect only non-human species such as bats (and palm civets and other mammals). SARS-CoV-2 is highly contagious in humans with the World Health Organization (WHO) designating it as a pandemic, the first ever caused by a coronavirus.

Pandemics such as the ongoing COVID-19 virus that leads to enormous loss of life globally can only be controlled by finding a vaccine or a very effective antiviral treatment. Finding a vaccine or specific antiviral treatment for such global pandemic of virus diseases requires rapid analysis, annotation and evaluation of metagenomic libraries to enable quick and efficient screening of nucleotide sequences. Traditional sequence alignment methods are not suitable since they are computationally intensive and cannot be easily scaled up as the number of sequences increase. Thus, there is a need for fast alignment-free techniques for sequence analysis [3, 4]. Further, one may have only short segments and/or incomplete fragments of nucleotide sequences to analyze [5]. Information theory and data compression algorithms provide a rich set of mathematical and algorithmic/computational tools to capture essential patterns in data that could be used for matching nucleotide sequences.

Genome sequences are inherently described by character strings and hence amenable to mathematical and computational techniques for extracting information. Exactly what information is being sought from such character strings depends on the string itself and the domain as well as the kind of application. Some targets of interest for analyzing genome sequences include:

- Various genes that constitute the genome.
- Identifying the origin of the genome sequence.
- Understanding the information content present in the coding and non-coding regions.
- Reconstructing the phylogenetic tree to study evolutionary patterns.

An important objective is to automate the above tasks so that a large number of sequences can be quickly, robustly and efficiently analyzed (as one of the steps in the endeavor for finding a vaccine).

A cursory glance at these character strings doesn’t tell us much about how they can be used for these applications. But a harmonious blending of complexity analysis with the field of information theory provides deep insight in this regard. Application of complexity measures on these information bearing character strings may reveal many surprising features that generally can’t be discerned by intuition or visual inspection of the data alone.

In this study, we propose a novel compression-complexity based distance measure for sequence analysis and identification. The paper is organized as follows. In section 2, a brief overview of genetic sequences and methods of analysis are described. Section 3 deals with the materials used (genome primary sequence data with their details) and the methods proposed in this study (novel distance measure for identification). Results on real data (nucleotide sequences) followed by their analysis and discussion can be found in section 4. We conclude with future research directions in section 5.

## 2 Genetic Sequences and their Analysis: An Overview

The basic building blocks of DNA and RNA are primary nucleobases, namely Cytosine (C), Guanine (G), Adenine (A), Thymine (T) and Uracil (U). A, C, G and T occur in DNA sequences and are known as DNA bases while A, C, G and U occur in RNA sequences and are called RNA bases. A string of these nucleobases forms a nucleic acid sequence that has the capacity to represent information. These information strings are called genetic sequences. Each species has unique characteristics differentiating it from other species and these characteristics are defined by the information content of the DNA sequences [6].

### 2.1 Genome and gene

The total DNA content (RNA for viruses) of an organism is known as the genome, thus representing the entire information coded in a cell, while a gene represents a section of the DNA that codes for RNA or protein. A genome consists of a sequence of multiple genes interspersed with non-coding sequences of nucleic bases [6, 7].

### 2.2 Genome sequence comparison

Genome data classification comes under the broad field of bioinformatics, an established multidisciplinary field for over three decades, encompassing physical and life sciences, computer science and engineering. Many fundamental problems in the fields of medicine and biology are being tackled using the tools of bioinformatics. The main requirement for accomplishing such tasks is the availability of sequenced genome data. This has been the focus of researchers for the past few decades and efforts have been put by the National Institutes of Health (NIH) to establish Genbank^®2^, a genetic sequence database containing annotated collection of all publicly available DNA sequences. Ever since its inception in 1982, there has been an exponential rise in the number of sequences in Genbank. This has provided the required resources for researchers and industry people alike for delving in to the field of bioinformatics.

Among the various aspects involved in bioinformatics, one key element is sequence comparison or analysis of sequence similarity [8, 9]. This is used in database searching, sequence identification and classification, phylogenetic tree^3^ creation and in gene annotation and evolutionary modeling. Since it is impossible to recreate/simulate past evolutionary events, computational and statistical methods for comparison of nucleotide and protein sequences are used for these kinds of studies [10, 11].

There are basically two kinds of sequence comparison methods:

- Alignment based methods: These involve either shifting or insertion of gaps in sequences for alignment of two or more sequences, which make these methods computationally intensive.
- Alignment-free methods: These are computationally less intensive methods that consider the genome sequences as character strings and use distance-based methods involving frequency and distribution of bases [12–14]. Our focus in this paper is on alignment-free methodology, especially on using complexity measures for sequence comparisons.

Sequence comparison and genome data classification got a boost in the early 1990s with the use of data compression algorithms that have the ability to identify regularities in sequences [15]. They provided a means to define distances between two sequences that greatly aided in the comparison of sequences. The history behind the usage of data compression algorithms in this field has been elucidated by Otu *et al.* in [15]. We succinctly summarize that history here.

The first attempt at using data compression for phylogenetic tree construction was by Grumbach *et al.* in [16]. They explored the idea of compressing a sequence *S* using a sequence *Q*, where the degree of compression obtained by doing so would be an indicator of the distance between them. Although their definition was not mathematically valid, it set a platform for researchers to explore in this area. Varre *et al.* [17] defined a transformation distance when sequence *Q* is transformed to sequence *S* by various mutations like segment-copy, segment-reverse copy and segment-insertion. Li *et al.* [18] define a relative distance measure by using a compression algorithm called GenCompress [19] that is based on approximate repeats in DNA sequences. Using the concept of Kolmogorov complexity, the compression algorithm has been used to propose a distance between sequences *S* and *Q*. But Kolmogorov complexity, [20] being an algorithmic measure of information and a theoretical limit, can’t be directly computed but only approximately estimated [21]. Hence it is not an optimum choice as a complexity measure.

Even though the idea of relative distance is an efficient one, GenCompress is a complicated algorithm that is computationally intensive. To overcome the above mentioned difficulties, Otu *et al.* [15] proposed similar but computationally simpler relative distance measures based on the Lempel-Ziv (LZ) [22] complexity measure. Given two sequences *S* and *Q*, sequences *SQ* and *QS* are formed by concatenation^4^. These four sequences are used to define four distance measures using the LZ complexity measure, as given below:

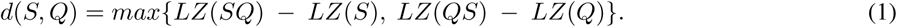

To eliminate the effect of length of the sequence, a normalized measure is defined as follows:

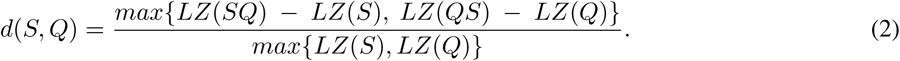

A third distance metric based on *sum distance* is defined as follows:

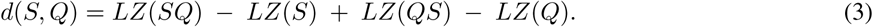

Finally, normalized version of the sum distance is defined as:

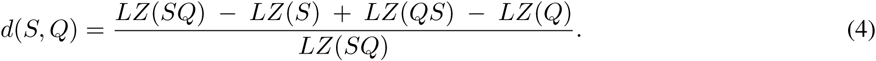

Using these distance measures on mtDNA (mitochondrial DNA) samples of a wide range of eutherans (placental mammals), they have successfully re-created phylogenetic trees showing the evolutionary patterns. Other researchers have used these and slight variants of these measures to identify families of coronaviruses, mammals, vertebrates and salmons. Interested readers are referred to [23–30] for further details on these. Apart from these complexity based measures, distance measures using Markov chain models [31–33] and measures of probability [34–36] have also been proposed for the study of genome identification.

### 2.3 Effect of data length on complexity of sequences

Monge *et al.* in [29] have pointed out that complexity is not uniform throughout a genome. Regions including genes are more regular and have less complexity than regions that don’t include a gene. This raises the issue of the length of the genome to be analyzed. Since the complexity is not uniform, it will be inaccurate to use the entire genome sequence for analysis and may possibly give erroneous results. Also the use of complete genome/gene sequences is computationally intensive and is practically infeasible for scaling up for matching a large number of genetic sequences. [5].

Our primary interest in this work lies in showing that it is not necessary to have the entire genome/gene for complexity analysis. In this work, we use LZ and Effort-To-Compress (ETC) [37] complexity measures to analyze short length segments that are randomly chosen from genome sequences. In particular, short length contiguous segments (length < 100) are randomly chosen from the sequence for analysis.

## 3 Materials and Methods

In this section, we describe in detail the data that was used in this study as well as the various methods that were employed for automatic identification of sequences.

### 3.1 Sequence analysis of coronaviruses (SARS-CoV-1)

We first analyzed the genome primary sequences of the following viruses: SARS coronavirus Urbani (AY278741.1), SARS coronavirus BJ01 (AY278488.2) and Avian Infectious Bronchitis coronavirus reference sequence (NC_001451.1). The first two virsues belong to the SARS-CoV-1 strain. The genome sequences were obtained from Genbank database, the details of which were mentioned in Section 2.2. Table 1 gives the details of the genome sequences that we use for the analysis.

**Table 1:** Genbank accession number, name, abbreviation and length of coronaviruses used for analysis.

| S.No | Accession Number | Genome | Abbreviation | Length |
| --- | --- | --- | --- | --- |
| 1 | AY278741.1 | SARS coronavirus Urbani | Urbani | 29727 |
| 2 | AY278488.2 | SARS coronavirus BJ01 | BJ01 | 29725 |
| 3 | NC_001451.1 | Avian infectious bronchitis coronavirus reference sequence | IBV | 27608 |

#### 3.1.1 0-1 sequences

The first step in our analysis is the conversion of primary sequences into 0-1 sequences before we can evaluate complexity values. For this, we have considered three different methods to categorize the nucleotide bases and map them to the symbols 0 and 1 based on their:

- Chemical structure
- Carboxylic acid group they belong to
- Strength of the hydrogen bonds

The four bases are mapped into two classes (labelled 0 and 1) in order to create a 0-1 sequence. For every input primary sequence, we create three independent sets of 0-1 sequences using three different methods as described in Table 2. It has been shown in [38] that these three characteristic sequences give the complete information of the primary sequence.

**Table 2:** Mapping of DNA bases in to three independent sets of 0-1 sequences using three different methods.

| Set No. | Method based on | Bases mapped to 0 | Bases mapped to 1 |
| --- | --- | --- | --- |
| 1 | Chemical Structure | Purine- {A,G} | Pyrimidine {C, T} |
| 2 | Carboxylic acid group | Amino- {A,C} | Keto- {G,T} |
| 3 | Strength of hydrogen bond | Weak H-bonds {A,T} | Strong h-bond {G,C} |

#### 3.1.2 Lempel-Ziv (LZ) and Effort-To-Compress (ETC) complexity measures

For measuring complexity of short length segments of the nucleotide sequences, we have used Lempel-Ziv (LZ [22]) and Effort-To-Compress (ETC [37]) complexity measures. Lempel–Ziv complexity (LZ), a popular and widely used complexity measure, estimates the degree of compressibility of an input sequence. Effort-To-Compress, a more recently proposed complexity measure (by our research group), determines the number of steps required by the Non-Sequential Recursive Pair Substitution Algorithm to compress the input sequence to a constant sequence (or a sequence of zero entropy). It should be noted that both LZ and ETC are complexity measures derived from lossless data compression algorithms (hence we term them as compression-complexity measures). It has been demonstrated that both LZ and ETC outperform Shannon Entropy in characterizing complexity of noisy time series of short length arising out of stochastic (markov) and chaotic systems [37, 39, 40]. Further, ETC consistently performs better than LZ in a number of applications as shown in recently published literature [39–42]. For details of how to compute LZ and ETC on actual input sequences, we refer the readers to [22, 37, 43].

#### 3.1.3 Distance measure and identification

We propose a very simple criteria for identification of sequences by proposing a distance measure which is computed using a compression-complexity measure (LZ or ETC). Let us say that we have genome sequences of three viruses *V*_1_, *V*_2_ and *V*_3_. Firstly, we form new sequences *V*_1_*V*_2_ and *V*_2_*V*_1_ by concatenation^5^. We then compute the complexity measures *ETC*(*V*_1_), *ETC*(*V*_2_), *ETC*(*V*_1_*V*_2_) and *ETC*(*V*_2_*V*_1_) (similarly for LZ). In line with what has been used by Otu *et al.* [15] in Equation 4, we propose a distance measure given by the average of the relative distances between the complexity values of the two concatenated sequences *V*_1_*V*_2_ and *V*_2_*V*_1_. Mathematically, they are described as:

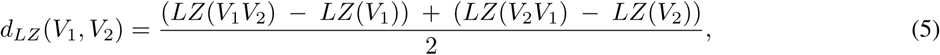

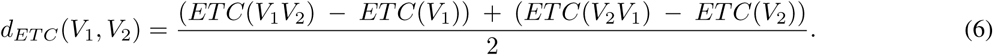

Note that the above distances will always be non-negative and symmetric^6^. In a similar fashion, we determine the distances *d*_*ETC*_ (*V*_1_, *V*_3_) (*d*_*LZ*_(*V*_1_, *V*_3_)) and *d*_*ETC*_ (*V*_2_, *V*_3_) (*d*_*LZ*_(*V*_2_, *V*_3_)). We then determine the minimum of the set {*d*(*V*_1_, *V*_2_), *d*(*V*_1_, *V*_3_), *d*(*V*_2_, *V*_3_)}^7^. We identify those viruses *V*_*i*_ and *V*_*j*_ which have the minimum distance to belong to the same group.

In our experiment, the sequences are converted to three independent sets of 0-1 sequences (by using the three methods listed in Table 2) and the LZ and ETC distances are independently calculated for all three sets. The average values of these are taken as the distance between the two sequences. In order to automatically identify the SARS viruses, the two SARS coronaviruses (Urbani and BJ01) should have the minimum distance (in complexities) as compared with the distance between the avian strain and any of the SARS coronaviruses (Urbani and BJ01). This method of identification can be easily extended if there are more than 3 sequences.

## 4 Results and Discussion

Figure 1 depicts the boxplots of pairwise distances (for LZ and ETC based measures) between the three viruses – Avian, BJ01 and Urbani, estimated for 100 short contiguous segments of length 30 nucleotide bases each of which were chosen independently at random locations of the entire genome sequence.

**Figure 1:**
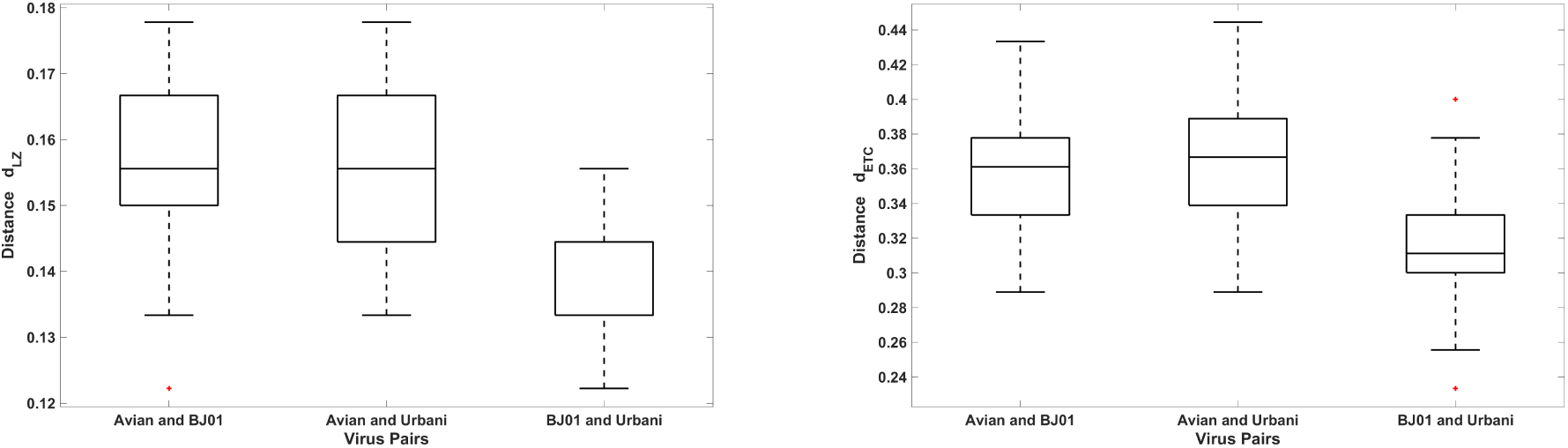
Boxplots of pairwise distance values for the three viruses – Avian, BJ01 and Urbani, calculated for 100 short contiguous segments of length 30 nucleotide bases chosen independently at random locations of the entire sequence. Left: *d*_*LZ*_, Right: *d*_*ETC*_.

The mean pairwise distances (and standard deviations) are reported in Table 3. As it can be seen, both LZ and ETC based distance measures yield the least value for the SARS-CoV-1 virus pair.

**Table 3:**
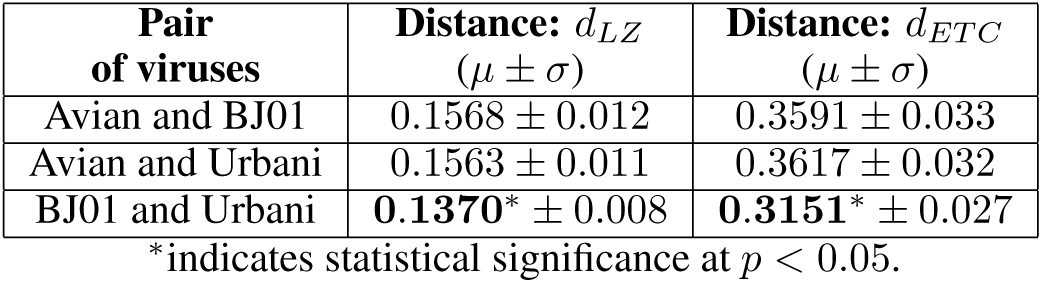
Pairwise mean distances for the three viruses – Avian, BJ01 and Urbani. We have averaged across 100 short contiguous segments of length 30 nucleotide bases each. These 100 were chosen independently at random locations of the entire genome. The mean distance between the two SARS-CoV-1 viruses BJ01 and Urbani is the least for both LZ and ETC measures.

The results were statistically validated with 95% confidence interval plots as shown in Figure 2. Based on the sample data, at an overall error rate of 5%, we can conclude that both LZ and ETC are able to identify the SARS coronaviruses from the given set of viruses using only short contiguous segments consisting of 30 nucleic bases chosen independently from random locations (100 such segments) of the entire sequence.

**Figure 2:**
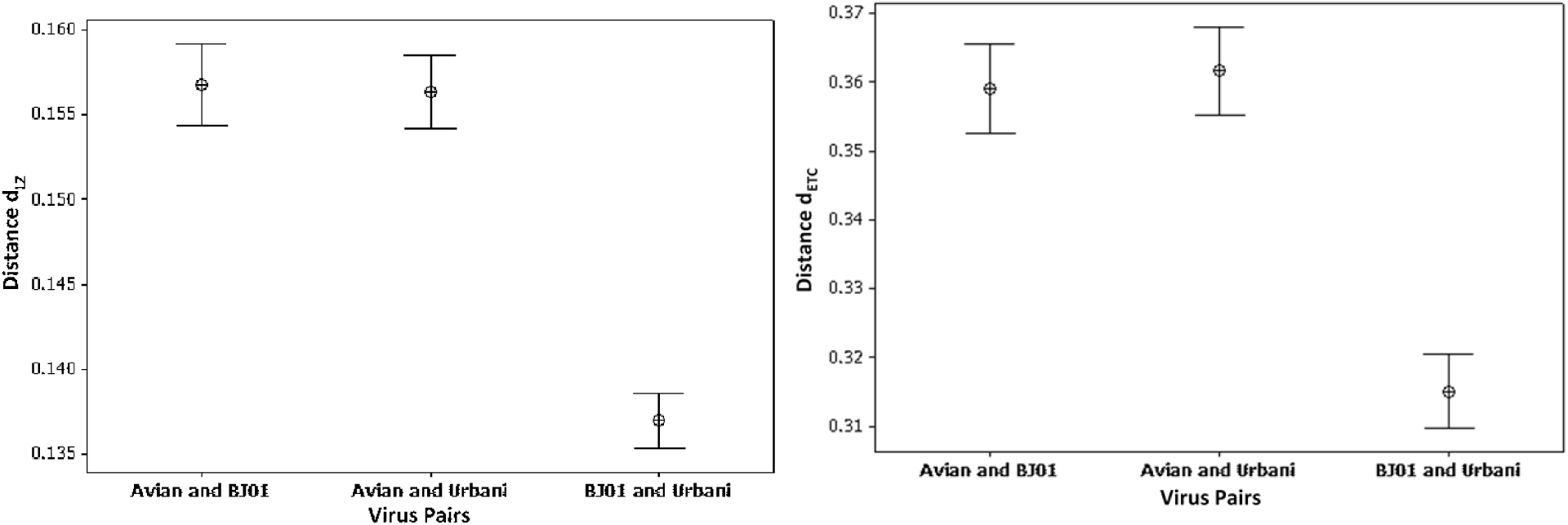
95% confidence interval for mean *d*_*LZ*_ (left) and *d*_*ETC*_ (right) distance measures for the three viruses (pairwise) – Avian, BJ01 and Urbani. The distance between the two SARS-CoV-1 viruses is clearly the least for both LZ and ETC.

### 4.1 Distinguishing SARS-CoV-1 vs. SARS-CoV-2

Having demonstrated the efficiency of LZ and ETC based distance measures in successfully distinguishing viruses by analyzing very short segments of nucleotide sequences, we extend our work to identify SARS-CoV-2 virus from SARS-CoV-1 viruses. To this end, we use the following sequences (Table 4) for this experiment.

**Table 4:** Genbank accession number, name, abbreviation and length of coronaviruses used for analysis.

| S.No | Accession Number | Genome | Abbreviation | Length |
| --- | --- | --- | --- | --- |
| 1 | AY278741 | SARS-CoV-1: Urbani | Urbani | 29727 |
| 2 | AY278488 | SARS-CoV-1: BJ01 | BJ01 | 29725 |
| 3 | NC_004718.3 | SARS-CoV-2: Reference Sequence | SARS-CoV-2 | 29751 |

Figure 3 depicts the boxplots of pairwise distances (for LZ and ETC based measures) between the three viruses – SARS-CoV-1:BJ01, SARS-CoV-1:Urbani and SARS-CoV-2 estimated for 300 short contiguous segments of length 25 nucleotide bases each of which were chosen independently at random locations of the entire genome sequence. The mean pairwise distances (and standard deviations) are reported in Table 5. It was found that only ETC based distance measure yielded the least value (statistically significant) for the SARS-CoV-1 virus pair (BJ01 and Urbani), and not the LZ based distance measure. The statistical validation of the results is depicted using 95% confidence interval plots as shown in Figure 4. Based on the sample data, at an overall error rate of 5%, we can conclude that ETC is able to distinguish between SARS-CoV-1 and SARS-CoV-2 viruses by using only short contiguous segments consisting of 25 nucleic bases chosen independently from random locations (300 such segments) of the entire sequence. LZ based distance measure fails to do so.

**Table 5:** Pairwise mean distances for the three viruses – SARS-CoV-2, BJ01 and Urbani. We have averaged across 300 short contiguous segments of length 25 nucleotide bases each. These 300 segments were chosen independently at random locations of the entire genome. The mean distance between the two SARS-CoV-1 viruses BJ01 and Urbani is the least for both LZ and ETC measures. This result is statistically significant only for the ETC based measure.

| Pair of viruses | Distance: $d_{LZ}$<br>($\mu \pm \sigma$) | Distance: $d_{ETC}$<br>($\mu \pm \sigma$) |
| --- | --- | --- |
| SARS-CoV-2 and BJ01 | $0.1648 \pm 0.014$ | $0.3801 \pm 0.033$ |
| SARS-CoV-2 and Urbani | $0.1625 \pm 0.015$ | $0.3749 \pm 0.038$ |
| BJ01 and Urbani | $0.1608 \pm 0.014$ | **$0.3661^* \pm 0.032$** |

\*indicates statistical significance at $p < 0.05$ .

**Figure 3:**
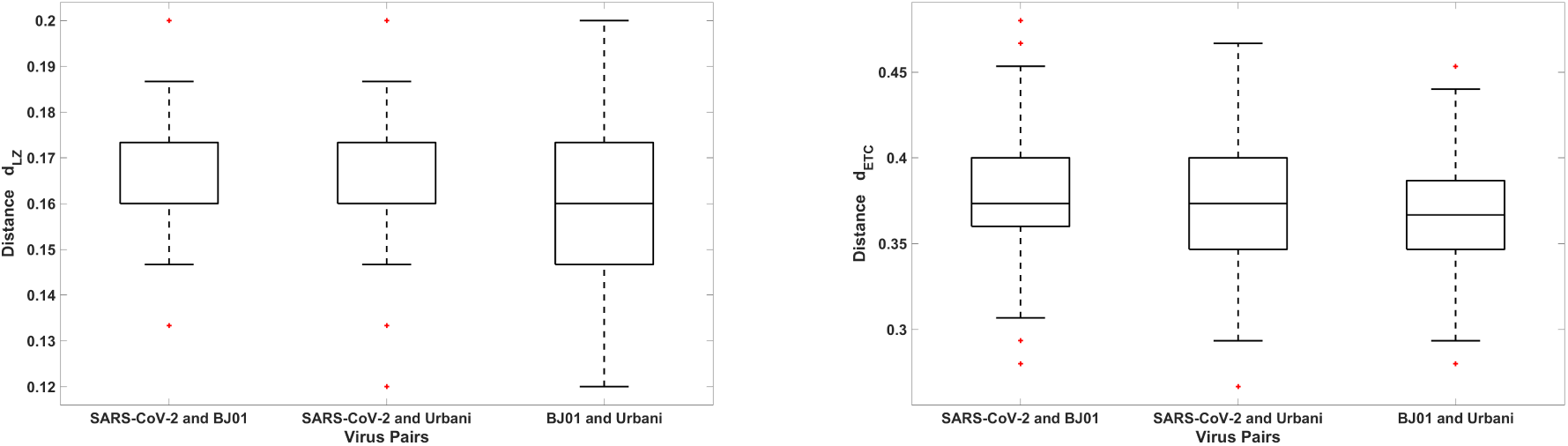
Boxplots of pairwise distance values for the three viruses – SARS-CoV-2, BJ01 and Urbani calculated for 300 short contiguous segments of length 25 nucleotide bases chosen independently at random locations of the entire sequence. Left: *d*_*LZ*_, Right: *d*_*ETC*_.

**Figure 4:**
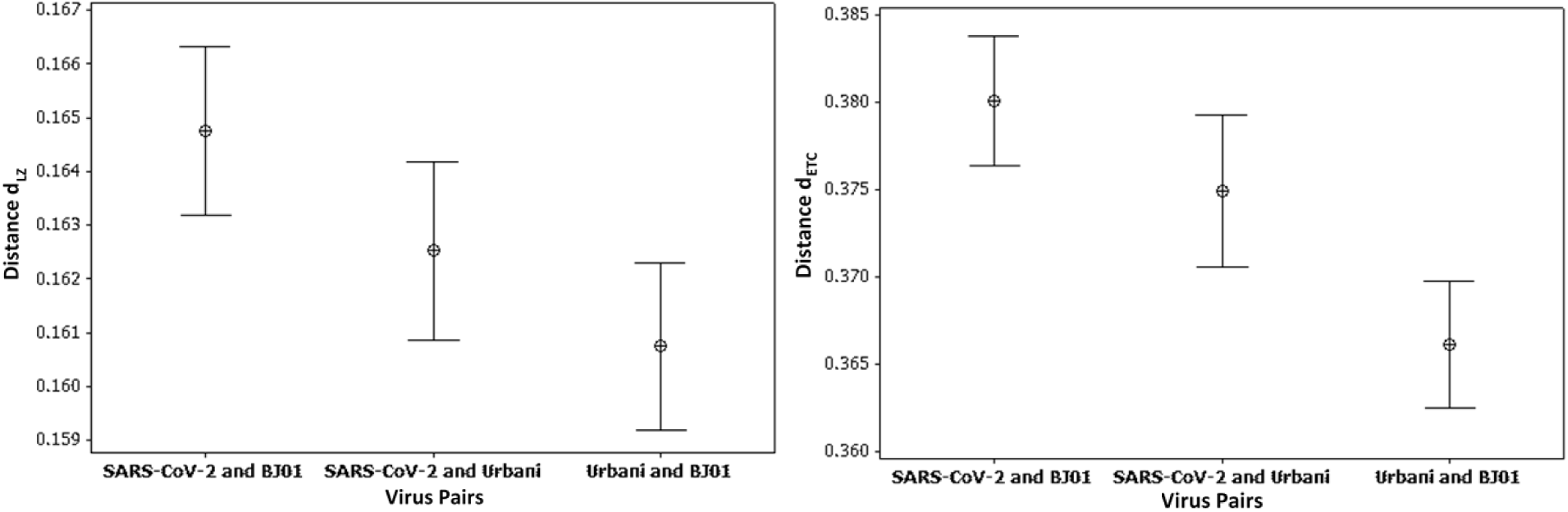
95% confidence interval for mean *d*_*LZ*_ (left) and *d*_*ETC*_ (right) distance measures for the three viruses (pairwise) – SARS-CoV-2, BJ01 and Urbani. The distance between the two SARS-CoV-1 viruses is clearly the least for ETC, but not for LZ.

To highlight the effect of segment length on distinguishing SARS-CoV-1 virsues from SARS-CoV-2 virus, we plot the pairwise distances (both LZ and ETC measures) for the three viruses for a *single* randomly chosen contiguous segment of length 5000 bases in Figure 5(left) and for another *single* randomly chosen contiguous segment of length 25 bases in Figure 5(right). It is evident that only ETC based measure is able to yield the least distance between the two SARS-CoV-1 pair of viruses for both lengths. LZ based distance measure fails to identify this pair for the short length segment and instead yields the least distance for the pair SARS-CoV-2 and Urbani which is not desirable.

**Figure 5:**
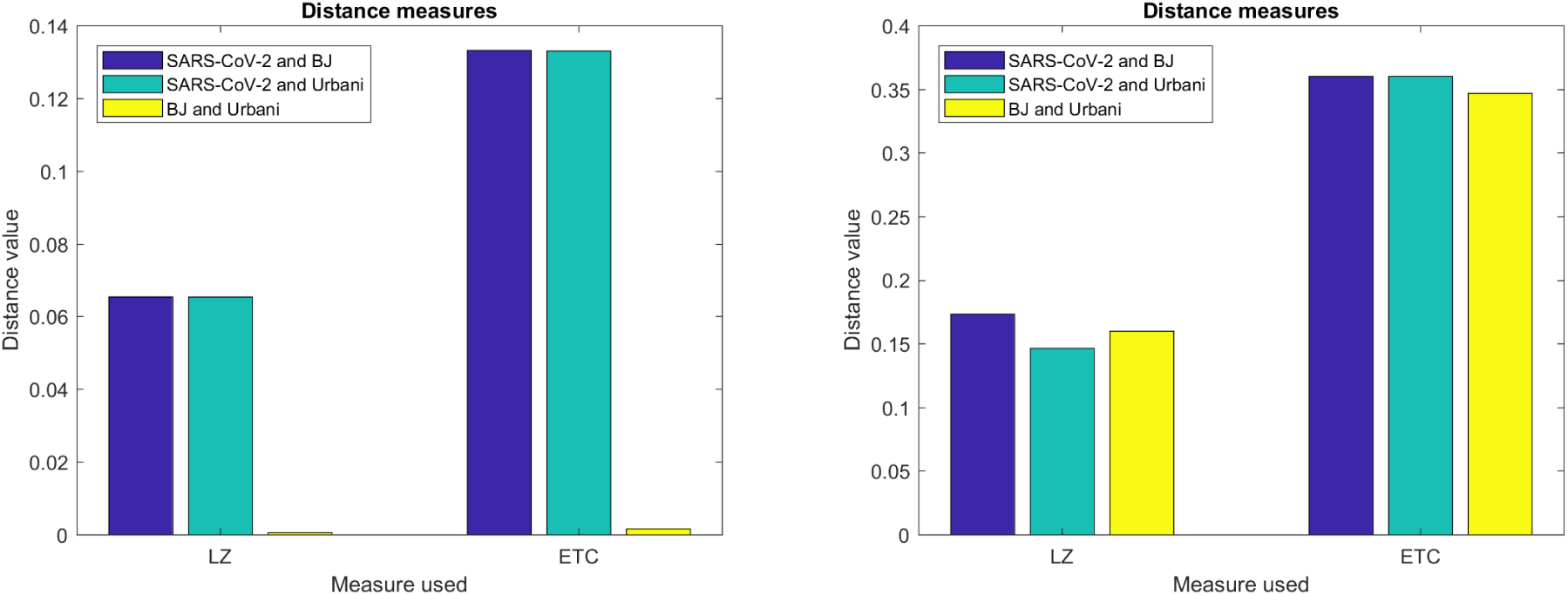
Effect of segment length on identification. Pairwise distances for the three viruses – SARS-CoV-2, BJ01 and Urbani computed using ETC and LZ based distance measures for a *single* randomly chosen contiguous segment of length 5000 bases (left) and 25 bases (right) from the nucleotide sequences. Only ETC correctly yields the least distance pair for the 2 SARS-CoV-1 viruses (BJ01 and Urbani) for both lengths.

Though these are still preliminary results, they are highly encouraging to further test ETC based distance measure for automatic identification/segregation of nucleotide sequences using only short segments.

## 5 Conclusion and Future Research Directions

Compression-complexity measures such as LZ and ETC which are based on lossless compression algorithms are good candidates for developing fast alignment-free methods for genome sequence analysis, comparison and identification. The main reason for this is their ability to characterize and analyze information in biological sequences with very short length contiguous segments. As we have demonstrated in this study, our preliminary results suggests that ETC could be very useful for identifying an unknown sequence from a large database of nucleotide sequences since we can quickly compute the measure on the candidate sequences for a small set of nucleic bases. LZ complexity requires slightly larger nucleotide sequences and that needs more computation. Other information theoretic methods in literature which employ Shannon Entropy, Mutual Information etc. would also need larger nucleotide sequences for computation and are not robust to noise. Some areas for further research are:

1. We have presented only preliminary results in this study and there is a need to rigorously test on distinguishing more sequences to further establish the reliability, robustness and universality of the proposed approach.
2. Construct a complete phylogenetic tree using the distance measure that we have proposed.
3. Compare ETC and LZ based distance measures with other methods in literature – both alignment-based and alignment-free methods.
4. Integrate ETC with existing open source packages that perform genetic sequence comparison and matching. To this end, we provide an open MATLAB^®^ and Python implementation of ETC that can be freely downloaded and used (link provided below).

The ideas presented in this study could potentially be extended further to enable medical practitioners to rapidly and automatically identify an unknown coronavirus sample (to be either a SARS coronavirus strain or a non-SARS coronavirus) in a fast and efficient fashion using only short and/or incomplete segments of genetic sequences. Further speed up can be obtained by parallelizing the analysis on individual short segments. Potentially, the need for sequence assembly can be completely circumvented.

## Software implementation of ETC

We provide open implementation of ETC (in MATLAB^®^ and Python) for free download and use (for research and academic purposes only). Please visit: https://sites.google.com/site/nithinnagaraj2/journal/etc.

## Acknowledgements

The authors would like to thank Gayathri R Prabhu (Indian Institute of Technology, Chennai) for helping with some of the simulations. NN would like to thank Pranay S Yadav (National Institute of Advanced Studies, Bengaluru) for the Python implementation of ETC and for useful discussions and suggestions.

## Footnotes

1 SARS-CoV-1 and SARS-CoV-2.

2 http://www.ncbi.nlm.nih.gov/genbank

3 A phylogenetic tree, also called an evolutionary tree, is a tree diagram that shows the evolutionary relationships among different species according to the composition of their genes.

4 *Q* is appended at the end of sequence *S* to yield the new sequence *SQ*.

5 *AB* is the new sequence obtained by simply concatenating sequence *B* at the end of sequence *A*.

6 *d*(*A, B*) *≥* 0, *d*(*A, A*) = 0 and *d*(*A, B*) = *d*(*B, A*). The triangle inequality is also likely to hold.

7 Here *d*(*·, ·*) could be either *d*_*ETC*_ or *d*_*LZ*_.

